# Group size dependent selection for cooperation versus freeloading in collective chemical defence

**DOI:** 10.1101/2025.09.03.672859

**Authors:** Sophie Van Meyel, Raphael Ritter, Heikki Helanterä, Carita Lindstedt

## Abstract

Individuals can cooperate in groups to achieve greater fitness, but the contribution of each individual to the collective good may depend on group size and on strategies employed by other members. How environmental contexts influence selection for cooperation is currently poorly understood and lacks experimental evidence from systems exhibiting behavioural plasticity in collective acts. This study investigates group size dependent selection on collective chemical defence in a gregarious pine sawfly (*Neodiprion sertifer*). Larvae perform chemical defence by secreting a costly deterrent fluid while adopting a U-posture. By manipulating group size and the individual’s ability to deploy the defensive fluid for the collective good, we show that survival against predation is higher in cooperative groups and that the benefits gained via collective defence are more pronounced in smaller groups. As predicted by theory, individuals participate less in collective defence in larger groups than in smaller ones. This lower contribution is not attributed to higher life-history costs from increased resource competition. Altogether, these results suggest that selection for cooperation in public goods is group size dependent, promoting cooperation in smaller groups whereas the relative fitness of freeloaders is higher in larger groups.

---

Evolution of group living and cooperation within a group has played a pivotal role in the evolution of life. For example, in groups individuals can effectively acquire and share costly resources such as food^1^, enzymes^2^, or defensive compounds against predators and pathogens^3,4^ known as *public goods*. Cooperation in public goods can evolve when sharing resources generates higher net benefits for an individual compared to what it can achieve alone^5,6^. For example, in *Pseudomonas aeruginosa* bacteria, defensive bacteriocin compounds against competing microorganisms are effective only when their concentration surpasses a critical threshold, requiring cooperation among multiple bacterial cells^4,7^. However, contributions to public goods are often facultative, allowing individuals the opportunity to adjust their investment to the prevailing ecological context^4,8^. Since the benefits of public goods are shared within the local group, individuals that do not incur the costs of cooperation (freeloaders) can still benefit from investment by other group members. Public goods systems are thus often vulnerable to freeloading which can corrupt cooperation^9,10^.

One area of interest in theoretical research has been the effect of group size on the evolution of cooperation^11,12^. Cooperation is expected to evolve more readily in smaller groups than in larger ones, particularly when the benefits of cooperation are shared among all group members^13,14^. In smaller groups, each individual receives a larger share of the collective benefits (1/n) and, as the group size increases, an individual’s contribution to public goods has a smaller impact on the overall outcome, while costs remain the same. As a result, in larger groups a cooperator receives a smaller share of the total benefits, reducing its pay-off from the cost of contributing^15^. Furthermore, if individuals are also in higher density in larger groups, closer interactions between freeloaders and cooperators may be facilitated, increasing the prevalence of exploitation ^15^. Cooperativeness can also decrease in larger groups if contribution to public goods becomes more costly at high densities^16^. This can happen if, for example, cooperation leads to more intensive resource competition in larger groups, thereby decreasing the levels of cooperation^17^.

Currently, most of the empirical evidence for the ‘Group size hypothesis’ comes from microbial systems, where both population density (e.g., through dilution) and the relative frequencies of genetic lineages of freeloaders and cooperators can be experimentally manipulated to assess the fitness of these two strategies across generations^15,18^. While density-dependent effects have been clearly demonstrated in microbial systems, these studies do not directly examine group size *per se,* and thus empirical evidence for its role in shaping facultative cooperative acts remains lacking. The lack of empirical evidence for the group size hypothesis has been partly due to the challenges associated with experimentally manipulating both the group size that individuals occur in and their level of contribution to cooperation. This issue is particularly evident in vertebrate and insect study systems, where facultative cooperation is more prevalent.

Here we solve this problem by using the gregarious pine sawflies (Hymenoptera: Diprionidae) as a study system. This system allows manipulation of both the group size and the level of cooperation and the quantification of the costs and benefits of public goods cooperation for individuals under different environmental contexts^8,19–21^. The larvae of this species live in dense groups ranging from 5 to over 100 individuals^20,22,23^, where they defend collectively against invertebrate and vertebrate predators (Fig. 1). When threatened by predators, these gregarious pine sawfly larvae display a defensive posture synchronously as a first line of defence, where they raise their head and posterior abdomen (U-posture). As a second, and more costly, stage of the defence they also simultaneously deploy a terpene and resin-rich fluid from their mouths^21,24^.

**Figure 1.**
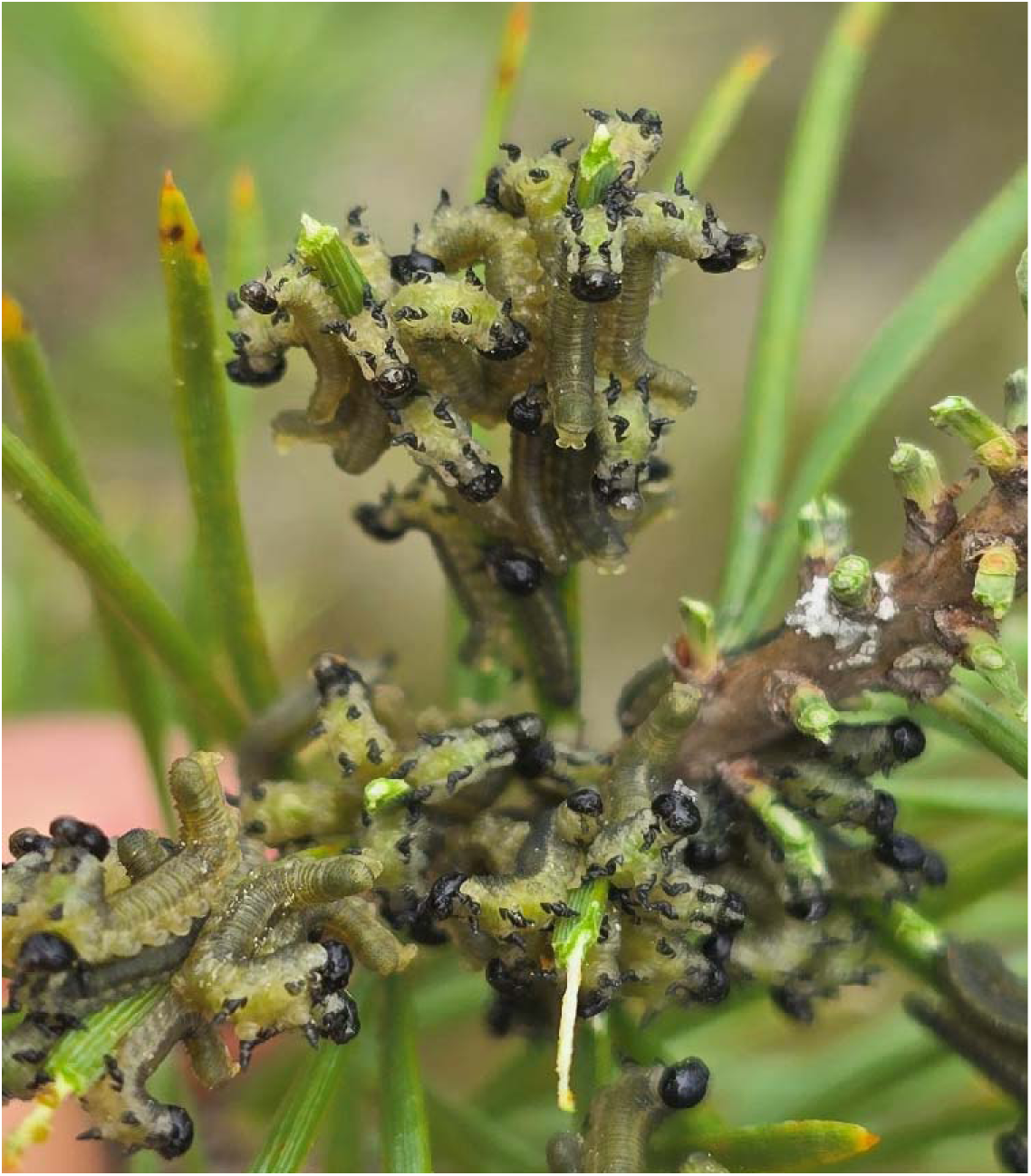
Group of defending larvae on a pine branch displaying their defensive behaviour in the wild. Some larvae display only a U-shaped defensive posture, while others combine it with the deployment of defensive fluid. Photo credit Raphael Ritter

In general, the responsive chemical defences such as those displayed by pine sawfly larvae, are considered to be public goods^25,26^. Prey individuals with responsive defences share the costs of educating predators to avoid prey with similar appearances in future encounters^25,27^. Additionally, chemically defended prey will benefit from being aggregated, as predators are less likely to attack prey individuals in groups where some or all group members are chemically defended^27,28^. Accordingly, pine sawfly larvae are unpalatable to both invertebrate and vertebrate predators^19,20,29–31^ and defensive fluid from multiple pine sawfly larvae forms a sticky barrier, hindering predators from capturing larvae without becoming coated in the fluid^24,30^. Larvae that have become depleted in defensive fluid have lower survival against predation^24^. Deploying and losing the fluid is thus costly for pine sawfly larvae, decreasing their growth as well as their future defence ability^8,22,32^. Larvae that deploy the fluid but do not lose it can reabsorb and reuse it. The deployment of defensive fluid is also a facultative trait^21^. Furthermore, some larvae consistently refrain from deploying fluid (so called ‘nil-individuals’) and benefit by growing faster; these individuals are suggested to be freeloaders^8^.

We used the above characteristics of *N. sertifer* to study experimentally (1) how group size and levels of cooperation, separately and in combination, influence survival against predators, (2) how contributions to collective chemical defence depend on group size, and (3) how group size affects the performance and life-history traits of individuals within groups. We predict that the survival benefits of cooperation should be more pronounced in smaller than in larger groups and, conversely, that the costs of freeloading should be lower in larger groups than in smaller groups. This is because defence may be less frequently required in larger groups because predation risk *per se* decreases as group size increases (dilution effect)^30,33^. Finally, if individuals decrease their contribution towards costly cooperation in larger groups, the proportion of individuals exhibiting defence should be lower in larger than in smaller groups. If the decrease in the level of cooperation in larger groups is due to increased resource competition, individuals should have a lower performance when in a large group compared to being in a small group.

## RESULTS

### Benefits of cooperation against predators are more pronounced in small groups than in large groups

The group size dependent benefits of cooperation were tested with a two-by-two full-factorial predation experiment in a seminatural setting using wood ants as predators. We manipulated both group size (5 and 20 larvae per group) and the level of cooperation (control and depleted). The manipulation of the level of cooperation was done by individually depleting the defensive fluid of larvae three times prior to the predation experiment in both group size treatments. In control groups, larvae sustained their defensive fluid.

Before exposing the larvae to ant predation, we first examined how defensive behaviour varied across treatment groups. Specifically, we analysed whether the probability of individuals displaying the defensive U-posture (with or without fluid) differed between group sizes and depletion treatments. We found that larvae in larger groups were less likely to display the defensive U-posture than those in smaller groups (LR-χ² = 20.46, df = 1, p < 0.0001). Furthermore, there were no significant differences between control and depletion treatment group in the probability of displaying the defensive U-posture (LR-χ² = 1.40, df = 1, p =0.24) and no significant interaction between group size and depletion treatment (LR-χ² = 0.0004, df = 1, p = 0.984). This indicates that group size influenced the probability of displaying the defensive U-posture independently of depletion treatment. However, when examining whether depletion treatment and group size affected the probability of deploying defensive fluid, we found that depleted larvae were significantly less likely to deploy fluid than control larvae (β = - 1.10, 95% ICr = [−1.58, −0.64], Fig. 2A). Additionally, larvae in larger groups were less likely to deploy fluid than those in smaller groups (β = −1.52, 95% ICr = [−1.97, −1.08], Fig. 2A). There was no significant interaction between group size and depletion treatment on the probability to deploy the fluid (β = −0.70, 95% ICr = [−1.67, 0.29]). This suggests that depletion treatment specifically reduced fluid (i.e., public good) deployment without affecting the overall defensive behaviour (U-posture) between depleted and control groups.

**Figure 2.**
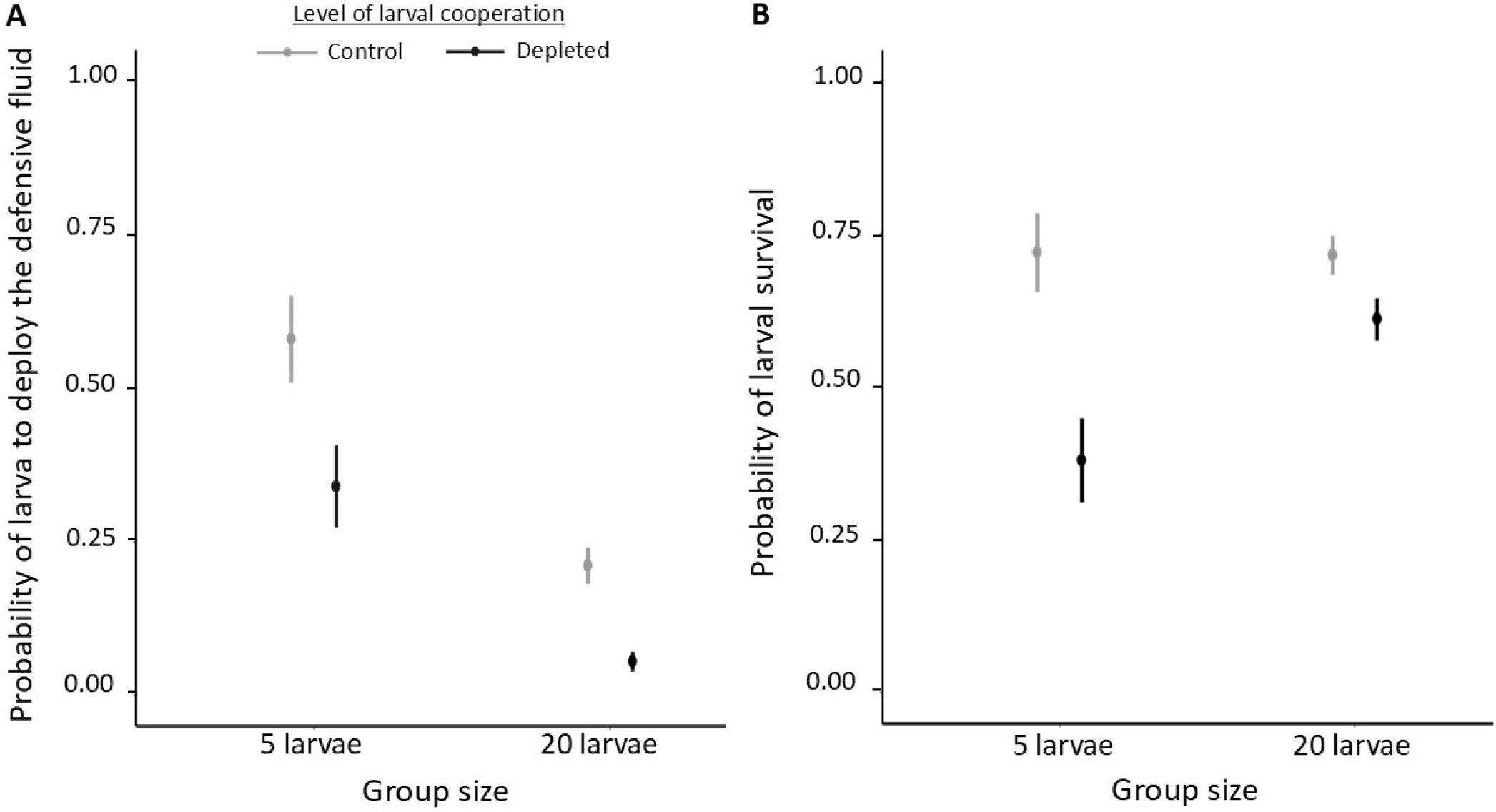
Effect of the group size (5 larvae or 20 larvae) and the level of cooperation (control or depleted) on collective defence and larval survival. (A) The probability of larvae deploying the defensive fluid is higher in small groups than in large groups (N =10 for both group sizes). Depleted larvae are less likely to deploy the defensive fluid than control larvae (N =10 for both treatments). (B) Depletion decreased the survival especially in the small groups (N =10 for all treatments).

When assessing individual survival against ant predation in relation to group size and the level of cooperation, we found a significant interaction between these two factors (LR-χ² = 14.01, df = 1, p = 0.0002, Fig. 2B). This indicates that the presence of non-defending individuals had a stronger negative impact on survival in smaller groups than in larger ones (Fig. 2B). In general, survival of individuals was lower in depleted groups in comparison to control groups (LR-χ² = 9.65, df = 1, p = 0.002), and in small groups in comparison to large groups (LR-χ² = 4.16, df = 1, p = 0.041).

### Individuals contribute less to the costly collective defence in larger than in smaller groups

To test whether individuals are more reluctant to contribute to collective defence in larger groups where benefit:cost ratio of cooperation is expected to be lower^15,26^, we conducted a factorial rearing experiment using a full-sib design under which *N. sertifer* larvae were reared in groups of 3, 10, or 20 larvae. We first measured the defensive behaviour of the individuals within the group by simulating an ‘attack’ on one randomly chosen larva per group (individual was poked once on the dorsal side with a capillary tube such as in)^8^. There were no significant differences in the probability of larvae displaying the defensive U-posture among group sizes (LR-χ² = 3.67, df = 2, p = 0.16, Fig. 3A). When using the absolute number of larvae per group adopting U-posture, we found that among larger groups more larvae displayed the U-posture (LR-χ² = 94.55, df = 2, p < 0.0001); group of 3 vs. 10 larvae (Z = −7.41, p < 0.0001), group of 3 vs. 20 larvae (Z = −9.68, p < 0.001), and group of 10 vs. 20 larvae (Z = −2.67, p = 0.02) (Fig. S1A).

**Figure 3.**
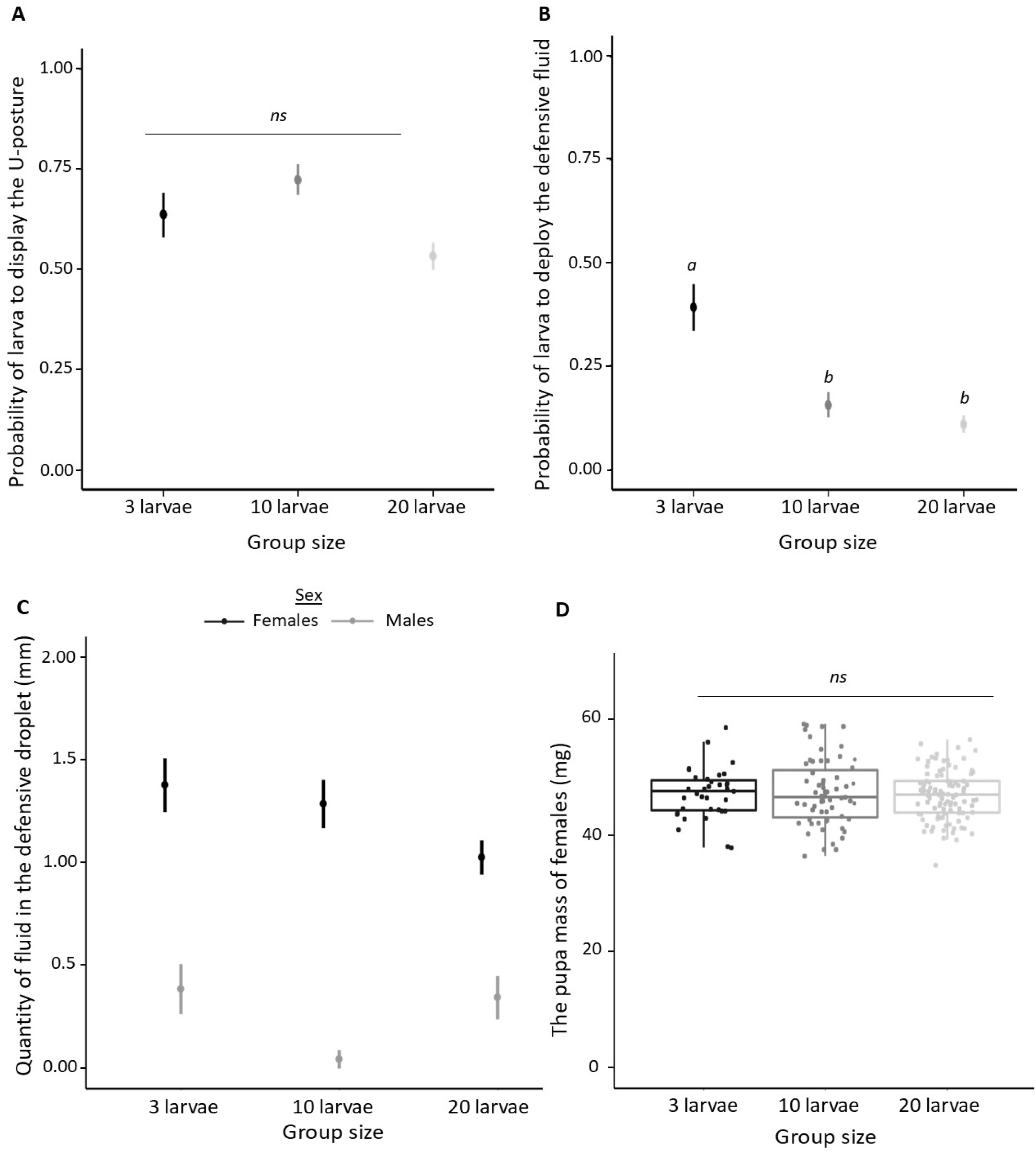
Effect of the group size on collective defence, individual defence and females’ pupal mass. (A) The probability of exhibiting the defensive U-posture after a simulated attack is similar in all the group size. (B) The probability of deploying defensive fluid after a simulated attack is higher in groups of 3 larvae than in bigger groups (10 or 20 larvae). (C) Females produce higher volume of the defensive fluid than males. Especially males from the group of 10 larvae produce lower volume in comparison to other group sizes. (D) Females’ have a similar pupal mass even when reared in different sized groups. The box plots represent the median and interquartile range, with whiskers extending to 1.5 times the interquartile range, each point represents one individual. ns = no significant difference between treatments. For pairwise comparisons, different letters correspond to p < 0.05.

The probability of deploying defensive fluid differed among group sizes (LR-χ² = 14.32, df = 2, p = 0.0008). More specifically, the larvae in larger groups (10 and 20) were less likely to contribute to collective defence than those in groups of 3 (group of 3 vs. 10 larvae: Z = 2.91, p = 0.01; group of 3 vs. 20 larvae: Z = 3.41, p = 0.002; group of 10 vs. 20 larvae: Z = 0.36, p = 0.93, Fig. 3B). However, the absolute number of larvae deploying the defensive fluid per group did not differ across group sizes (LR-χ² = 3.82, df = 2, p = 0.15, Fig. S1B) suggesting that the total number of larvae committing to costly collective defence behaviour remained constant.

Additionally, we assessed the contribution to chemical defence on individual-level by attacking each larva separately (i.e. larva defends itself). We found that the group size had a significant effect on the fluid deployment; individuals were less likely to deploy defensive fluid in groups of 10 larvae than in groups of 3 larvae (β = 1.51, 95% ICr [0.13, 3.37], Fig. S2A). Larvae in groups of 10 also tended to deploy less than those in groups of 20, but this difference was not significant (β = −0.41, 95% ICr [−1.85, 0.82], Fig. S2A). Furthermore, *N. sertifer* female larvae were more likely to deploy the defensive fluid than males (β = −1.25, 95% ICr = [−2.28, −0.28], Fig. S2B). When we analysed the effect of group size and sex on the volume of fluid deployed, there was an interaction between the group size treatment 10 and sex of the individual (β = −2.34, 95% ICr = [−4.34, −0.37], Fig. 3C). This indicates that especially in the group size of 10, males were deploying lower volumes of fluid in comparison to males from groups of 3 larvae and 20 larvae (β = 2.68, 95% ICr = [0.72, 4.46] and β = −2.63, 95% ICr = [−4.48, −0.93] respectively) or in comparison to volumes deployed by females under different group size treatments (Fig. 3C). Again, females deployed higher volumes of fluid than males (β = −1.47, 95% ICr = [−2.86, −0.07]).

### Life-history costs of contributing to the collective defence did not depend on the group size

The lower contribution to the collective defence in larger groups was not explained by increased life-history costs for an individual (Table 1): for both sexes, the developmental time, pupal mass or reproductive output did not depend on the group size treatment or individual’s contribution to collective defence (i.e. whether the larva deployed the defensive fluid during individual defensive measurements or not) or the interaction between these two factors (Table 1: both sexes, Fig. 3D, Fig. S3: females only).

**Table 1.** Effect of the group size, defensive behaviour and their interaction on larval performance of males and females separately.

| | Group size | | Defensive behaviour | | Group size × Defensive behaviour | |
| --- | --- | --- | --- | --- | --- | --- |
| | LR $\chi^2_2$ | P | LR $\chi^2_1$ | P | LR $\chi^2_2$ | P |
| <u>a) males' life history traits</u> |  |  |  |  |  |  |
| Larval developmental time | 1.85 | 0.40 | 0.28 | 0.60 | 3.78 | 0.15 |
| Pupal mass | 0.85 | 0.65 | 0.42 | 0.52 | 2.74 | 0.25 |
| <u>b) females' life history traits</u> |  |  |  |  |  |  |
| Larval developmental time | 4.15 | 0.13 | 0.04 | 0.84 | 0.04 | 0.98 |
| Pupal mass | 0.79 | 0.68 | 2.23 | 0.14 | 2.06 | 0.36 |
| Number of eggs produced | 2.59 | 0.27 | 1.52 | 0.22 | 0.0006 | 0.98 |

## DISCUSSION

Cooperation in public goods is expected to be more likely to evolve when group sizes are smaller due to increased net benefits and lower competition between group members^12,15,17^. Here we show that cooperation in collective chemical defence provides greater survival benefits in smaller groups while freeloading incurs lower survival costs in larger groups. We also show that individuals modified their contributions to collective defence accordingly, by contributing less to costly chemical defence when in large groups. As this was not explained by increased resource competition in larger groups but rather suggests that individuals adjusted their investment in costly collective defence due to group size dependent variation in its benefits.

One possible explanation for the higher benefits of defending in smaller groups is that the higher *per capita* risk of attack increases the direct benefits of defending. For example, displaying fluid can serve as a self-advertisement of an individual’s defensive capacity via visual, olfactory, or gustatory cues^34^ in addition to its function in simply ‘fighting’ against a predator^30,35–37^. Density-dependence in contribution to collective chemical defence was also supported by the behavioural data (Fig. 3A, B). While the proportion of larvae performing the defensive U-posture was similar across group sizes, the proportion of individuals participating in the more costly chemical stage of the defence decreased in larger group sizes. This suggest that collective defence in *N. sertifer* may continue to be effective as long as a critical number of individuals contribute in the vicinity of the attacked individual. In larger groups, this threshold can be reached with a smaller proportion of contributors, allowing more individuals to adopt a freeloading strategy without compromising their survival.

An alternative explanation for decreased cooperation in larger groups is the heightened associated costs; for example, due to increased competition for food^26^. While individuals were less likely to deploy defensive fluid in larger groups, we found no evidence that this behavioural strategy was driven by increased life-history costs. Larval development time, pupal mass, and female egg production were unaffected by group size even when we considered individual’s contribution to chemical defence in the last larval instar (i.e., if costs of being in a bigger group would only be visible in individuals that deployed and lost the fluid). This indicates that the energetic or physiological costs of defence did not differ across treatments. These results could be explained by our experimental conditions (i.e., food *ad libitum*) that may have minimized the potential costs. However, our field observations suggest that, at least in the early stages of outbreak, there are only one-to-two larval groups within each pine tree, suggesting that food competition is unlikely. During high density periods, food competition may however operate and our experimental design therefore reflects conditions typical of low population density.

We also found that similarly to other sexually dimorphic pine sawfly species^8,38^, the contribution to collective defence is sex-biased: female larvae were more likely to deploy the defensive fluid and higher volumes of it than males. This female-biased contribution was consistent across group sizes and was not associated with higher life-history costs (Table 1). In accordance with other studies (see also^8,39^), our study here suggests that female-biased investment in collective defence in immature life-stages is a consistent behaviour in several pine sawfly species. Based on the studies with eusocial haplodiploids, such female-biased investment can evolve if females are ancestrally pre-adapted for cooperative behaviour^40,41^. While *Neodiprion sertifer* does not exhibit advanced maternal care^21^, the size dimorphism between females and males could act as a pre-adapted trait for collective defence, whereby larger females may contribute more effectively to group protection against predator attacks (see ^42^ for sex-biased division of labour in termites driven by sexual size dimorphism). For example, it is possible that increased size allows females to simply store and produce more defensive fluid, or that they experience lower relative costs when deploying it^8^. The sex-biased investment in defence can also be promoted due to haplodiploidy and sex ratio adjustments it facilitates^43^. Pine sawfly populations are female-biased^44–47^ and ecological conditions can result in variation in the selection pressure for mother to produce the more cooperative sex (i.e., local resource enhancement)^48–50^. Since the reproductive value is lower for the individuals of the more common sex (“the rarer sex effect”)^51,52^, it may drive those individuals (in this case female pine sawfly larvae) to invest even more in cooperating, creating a positive feedback loop between sex-ratio bias and sex-biased investment in cooperation^43^.

The group size had a lower impact on the plasticity of the defensive response when individuals were defending themselves (i.e., larvae were attacked individually). In females, the contribution to defence showed a decreasing trend with increasing group size, similar to the pattern observed in group level measurements. However, with males, the pattern differed: contributions were lower in groups of 10 larvae compared to other group sizes (see Fig. 3C and Fig. S2B). This difference in the females’ and males’ responses to group size variation was more pronounced when we considered the volume of fluid deployed (see Fig. 3C). We do not have a clear explanation for this pattern. Other studies with pine sawflies suggest^53^ that individuals’ contributions to collective defence tend to decrease in male biased groups. However, in our experiment the sex-ratios did not differ between group size treatments (see Methods) suggesting that sex-ratio variation between treatments does not explain the observed pattern. Similarly, genetic variation was controlled across treatment groups using a split-family design, and both food availability and abiotic conditions were standardised between treatment groups. This suggests that the reduced male contribution to defence observed in groups of ten larvae may may be attributable to the specific social context of that group size treatment. Nevertheless, how the responses of female and male larvae to collective defence differ depending on the environmental context will provide an interesting avenue for future studies.

Behavioural plasticity enables individuals to maximize their fitness across varying environmental and social conditions. In public goods, such flexibility allows organisms to adjust their level of investment in response to environmental factors, such as group size, resource availability, or the other group members behaviour^26,54–56^. In *N. sertifer*, individual investment in chemical defence varies with group size in a manner consistent with group size dependent selection. Larvae exhibit a facultative response to the changing benefit:cost ratio associated with group size and per capita predation risk. This could be adaptive given that *N. sertifer* pine sawflies form regular outbreaks where social and ecological conditions can change fast for each generation. More generally, however, it remains unclear to what extent cooperative behaviour is genetically fixed or plastic due to environmental conditions. For instance, in *Pseudomonas aeruginosa*, the siderophore production is regulated by *pvd* genes, whose expression is plastically regulated in response to iron availability in the environment^57^. At the same time, mutations in these genes can give rise to ‘cheater’ genotypes, which do not contribute to the public goods and exploit siderophores produced by others^58,59^. In contrast, the precise genetic underpinnings of the contribution to cooperative behaviours in multicellular organisms, are much less understood. While positive but low heritability have been observed in wild meerkats^60^, wild long-tailed tits^61^, in cooperative breeding cichlids^62,63^ and in western bluebirds^64^, context-dependent variation in cooperation is prevalent^55,56^, and plasticity in traits is especially favoured in unpredictable environments^65^.

Finally, group size effects on benefit:cost ratio of cooperation have been suggested to play a critical role in the transition from simple group living to more complex cooperation within a group^66–69^. Accordingly, our results demonstrate that variation in group size shapes the benefit:cost ratio of cooperation among group members resulting in variation in the strength of selection toward cooperativeness. Our study design did not take into account how the group size interacts with the strength of kin-selection. However, the pine sawfly larvae exist mainly within full-sibling groups in nature suggesting that the strength of kin-selection toward cooperativeness does not vary extensively with the group size^53^. More generally, while the decreased predation risk in larger groups have shown to be a predominant explanation for the group living in primates, birds, fish and insects, our results contribute to the growing body of empirical evidence^17,69^ that suggests that predation risk and its interaction with group size can also be major ecological drivers of complex cooperation.

## SUPPLEMENTAL INFORMATION

The supplementary information includes three figures and their legends.

## ACKNOWLEDGEMENTS

This study was funded by the Academy of Finland via the projects no 257581 and 330578 (CL). We are grateful to Tuuli Salmi, Kaisa Suisto, Minna Salonen, Liina-Lyydia Jämsä and Venla Korhonen for helping with the maintenance of lab populations and collecting field data, and Ian C.W. Hardy for commenting on an early version of the manuscript.

## AUTHOR CONTRIBUTION STATEMENT

CL conceived the ideas and CL, RR and HH designed methodology; RR and CL collected the data; SVM and RR analysed the data; SVM led the writing of the manuscript, with input from RR and CL. All authors contributed critically to the drafts and writing and gave final approval for publication.

## DATA AVAILABILITY STATEMENT

On acceptance of this paper for publication, all data and final R-codes used in the analyses will be included as an open access database in DRYAD. For the review purposes all data and codes presented in figures and tables are included in a supplementary file.

## METHODS

### Effect of group size and levels of cooperation on larval survival against predators

In June 2023, 19 larval groups of *Neodiprion sertifer* were collected from Puumala (61.56082, 28.01005) in eastern Finland. Each group was at approximately the second instar stage. Larvae were fed *ad libitum* and maintained in standardized laboratory conditions (+ 20 C). When larvae were between 3^rd^ to 4^th^ instar, each initial group was split into two large groups of 20 larvae (N = 20) and two small groups of 5 larvae (N = 20) which were further randomly divided into the depleted and non-depleted treatments. Altogether, both depleted and non-depleted groups of 5 and 20 larvae were created, with N = 10 groups for each treatment, respectively.

During the depletion manipulation, larvae were gently pressed with a cotton swab on the dorsal side. When they deployed their defensive fluid, it was carefully removed with the swab. This was repeated twice on the following day. After the depletions, larvae had until the next morning to re-aggregate after the manipulations.

On the following day, the propensity of defence in response to a simulated attack by a predator was quantified before taking a group of larvae to the field. We randomly chose one larva from within each group and gently poked it once on the dorsal side with the tip of a pair of tweezers and scored whether or not it exhibited the U-posture, and whether or not defensive fluid was deployed.

Groups of larvae on their original pine twigs were transported into the field where the larval branches were attached to the bigger 40 cm long pine branches containing two smaller twigs that were installed next to the ant trail in the vicinity (2-5 m) of an ant nest. Altogether, we used 10 different mound building wood ant nests (*Formica s.str.*). Thus, we tested all four treatment groups at the same ants’ nest, ensuring that potential variation in activity or aggressiveness among ant nests was taken into account. The survival of larvae was recorded by counting the number of larvae within each group at the beginning of the experiment and again after seven hours.

#### Statistical analyses for the predation experiment

To confirm that the depletion treatment was successful, we first tested whether the probability of displaying the U-posture (with or without fluid) and the probability of deploying the defensive fluid differed between group sizes and depletion treatments. To analyse whether larvae raised their heads in a defensive U-posture, we fitted a generalized linear mixed-effects model (GLMM) with a binomial error distribution and a complementary log-log (cloglog) link function. We used this link due to the moderate imbalance between 0 and 1 outcomes in the response variable. Display of the U-posture (1 = yes, 0 = no) was the response variable and group size (5 larvae, 20 larvae), depletion treatment (control, depleted), and their interaction were used as fixed effects. The larval group ID was included as a random effect to consider potential influences of the common environment effects of larval group. Due to the singularity issues in the GLMM, where the variance of the random factor (larval group ID) was estimated as zero, we chose to use a Bayesian approach using the brms package (Stan) to analyse how a larva’s probability of deploying the defensive fluid differed between treatment groups. This allowed us to keep the larval group ID in the model and to stabilize estimation using weakly informative priors. A Bayesian approach was particularly suited here because it handles low group-level variation and imbalanced binary responses better than frequentist methods. We ran a Bayesian Generalized Linear Mixed Model (GLMM) with a Bernoulli distribution appropriate for binary (0/1 per larva) data and clog-log link function (see above), including the group size, depletion treatment and their interaction as fixed effects, and ID of the larval group as a random factor. We used weakly informative priors to regularize parameter estimation: normal (0,5) for fixed effects, student_t(3, 0, 10) for the intercept, and cauchy(0,5) for random effect standard deviations. The model was fitted using Hamiltonian Monte Carlo (HMC), with 4 chains of 4000 iterations.

To analyse the probability of larval survival against ant predation in different treatment groups, we used a generalized linear mixed model (GLMM) with binomial error distribution and a complementary log-log (cloglog) link function. Larval survival (1 = alive or 0 = dead) was used as the response variable and the depletion treatment and group size and their interactions were fitted as explanatory variables. Nest ID and larval group ID were included as random factors to consider potential variation in survival due to differences (e.g. activity, hunger level) among ant trails or ant nests.

### Effect of group size on the life-history traits and contribution to collective chemical defence

#### Neodiprion sertifer laboratory cultures

The experiment was conducted in Jyväskylä, Finland. All *N. sertifer* families descend from an outbred F1-generation of laboratory population established in 2016. This population was founded from individuals originating from 86 wild larval colonies collected from Central Finland (58 colonies, Puumala 62.066704, 27.351608 and 28 colonies, Pieksämäki 62.066704, 27.351608) and reared until adulthood in constant temperature (20 °C) and density (20 larvae per container) with fresh food (*Pinus sylvestris*) available *ad libitum*. Cocoons were individually stored into the Petri dishes that were kept in the same rearing conditions as larvae. After adults eclosed, females were allowed to mate (one male per female) and lay their eggs on a randomly chosen pine branches of a living tree (*P. sylvestris*) (Central Finland 60° 15’ 17,48” N, 26° 1’ 59,03” E). Each branch with a mated pair was covered with a mesh bag. Females oviposit eggs inside the pine needles and *N. sertifer* overwinters in egg stage^23^. Eggs die if the needle surrounding them dies^70^. Therefore, we had to use living trees and keep egg branches outdoors through the winter to allow the eggs to go through the obligatory diapause and develop.

In May 2017, the pine branches with eggs were collected and maintained in a constant temperature (+20 °C) until eggs hatched. Hatched larvae were fed *ad libitum* with pine branches, and fresh branches were provided twice per week. All larvae were reared in similar conditions under constant temperature (+20 ±2 °C) in a laboratory room that included both natural light coming from the windows (nights are very short in Finland during May and June) plus artificial light at all times, mimicking the normal central Finnish summer light conditions.

## Experimental design

Families with the minimum of 40 individuals were chosen for the experiment to be able to follow a full-sibling design. At the age of 10 days, 36 larvae per family were divided into three different group sizes: two groups of 3 larvae, one group of 10 larvae, and one group of 20 larvae. We chose to use two replicates of 3 larvae treatment per family to take into account the unbalanced number of individuals due to group size treatment. Larvae were kept in transparent plastic containers with fabric on top for ventilation in a condition described above. Altogether we had 16 families in the experiment from which 7 were offspring of parents collected from Puumala and 9 were offspring of parents collected from Pieksämäki. This resulted N = 27, N = 15 and N = 13 for larval groups in treatments of 3 larvae, 10 larvae and 20 larvae, respectively. All the treatment groups included similar amounts of females and males as there were no significant differences in the sex-ratios among group size treatments (LR-χ² = 0.39, df = 2, p = 0.825).

### Contribution to collective defence under different group size treatments

The group defence behaviour of larvae was measured on the third day of the experiment. We recorded the number of larvae that adopted the U-posture, and the number of these that deployed the defensive fluid, after one randomly chosen larva within the group was “attacked” by gently poking the larvae once with the tip of a pair of tweezers on its dorsum. We also measured the individual level contribution to the collective defence from the last larval instar at 16 days of age. For this part of the experiment, we used larvae from 14 families in total (N = 7 from Puumala and N = 7 from Pieksämäki). Each larva was individually gently pressed on the dorsal and ventral side with a capillary tube. We recorded whether or not the larva produced a defensive droplet. To measure the volume of any fluid produced, it was sucked into a 5uL capillary tube (Microcaps®, Drummond Scientific Co., Broomall, Penn.) the length of the liquid measured (using a digital calliper to the closest 0.1 mm) and converted to a volume. At the same time, we measured the body length of each larva using digital calliper to the closest 0.1 mm to control for the effect of body size on the amount of defence fluid produced.

### Effect of group size treatment on life-history costs

We recorded the development time of larvae to reach pupation (in days) and pupa weight (3-4 days after their pupation). To measure the reproductive output of females, females from different treatments were mated with males originating from the same treatment. Altogether, we had 31 pairs from groups of 3 larvae, 33 from groups of 10 larvae and 37 from groups of 20 larvae. Mating was allowed similarly to the P-generation (see above, *N. sertifer* laboratory culture). Mated pairs from different treatments were put on the same tree on the same day (one pair per each treatment per one host tree) to control for possible variation due to host plant quality. Similarly, mated pairs were taken outdoors to lay eggs in a similar weather condition (sunny and dry days, +18 to 25°C).

### Statistical analyses of the chemical defence and life-history traits

We used generalized mixed models to estimate how treatment affects how larvae contributed to common defence at the group level when one individual within the group was attacked. Display of the U-posture (yes/no) and deployment of fluid (yes/no) were treated as dichotomous variables modelled as binomial response variables with a cloglog-function. In all the models (both frequentist and Bayesian (see below), the population ID, family ID and group ID were included as random factors, to take into account any similarity due to common inheritances and environments. We also analysed the group-level defence using the absolute number of larvae per group that performed the U-posture and the number that deployed the defensive fluid (both are integer-valued count data). We used two generalised linear mixed-effect models (GLMM) with Poisson error distribution and log-link function. Each behaviour was modelled separately as the response variable, with group size (3, 10, or 20 larvae) as the only fixed effect. While individual-level models capture the probability of each larva performing the two defensive behaviours, absolute numbers reflect the total group-level mobilisation. A comparison of these two analysis methods allows us to distinguish between individual decision-making and potential collective thresholds in group defence.

To analyse how group size, sex and their interaction affects an individual’s fluid deployment and its volume (i.e. when individuals were directly attacked and defending on themselves), we used a Bayesian approach using the brms package (Stan) allowing us to keep the random factors in the model and to stabilize estimation using weakly informative priors (see above). We fitted a Bayesian Generalized Linear Mixed Model (GLMM) with a Bernoulli distribution and cloglog link function. Deployment of the defensive fluid (1 = yes, 0 = no) was the response variable and group size (3 larvae, 10 larvae, 20 larvae), sex (female, male), and their interaction were entered as fixed effects. Population ID, family ID and group ID were included as random factors, to take into account any similarity due to common inheritances and environments. Weakly informative priors were used: normal (0,5) for fixed effects, student_t(3, 0, 10) for the intercept, and cauchy(0,5) for random effect standard deviations. The model was fitted using Hamiltonian Monte Carlo (HMC), with 4 chains of 4000 iterations^38,71^. For the volume of fluid, the same fixed and random effect’s structure and priors were used as above. We fitted a Bayesian GLMM with a gamma error distribution and a log link function since the data were positively skewed. Because the gamma distribution does not allow zeros, and measurement precision was 0.01, we used “measured volume + 0.01” as the response variable^8^.

Due to sexual dimorphism, life-history traits were also analysed separately for females and males^8^. We fitted three generalized linear mixed-effects models (LMERs) with development time, pupa mass, and egg production (the latter only for females) as response variables. To test the effect of the defensive behaviour (1 = yes or 0 = no) on individual’s performance, we included the defensive behaviour as a fixed factor together with the group size and their interactions.

All statistical analyses were performed with the software R v4.4.1 (http://www.r-project.org/) loaded with the package *car* ^72^, *lme4*^73^; *ggplot2*^74^; *DHARMa*^75^, *emmeans*^76^*, brms*^77^. The residuals from each model were visually examined for deviations from normality and for heteroscedasticity^78^, using the testResiduals() function from the DHARMa package, which performs standard diagnostic checks based on simulated residuals^75^. In Bayesian models, convergence was evaluated by ensuring Rhat values were below 1.01 for all parameters and effective sample sizes (ESS) were sufficient, with no divergent transitions detected. Posterior predictive checks confirmed good model fit^77^.

When an interaction was not significant, we removed it and refitted a simplified model without the interaction term. For models where an interaction was significant, we performed pairwise comparisons between treatments using estimated marginal means (emmeans) and applied Tukey corrections for multiple comparisons^79^.

