## Supplementary material for "Group size dependent selection for cooperation versus freeloading in collective chemical defence"

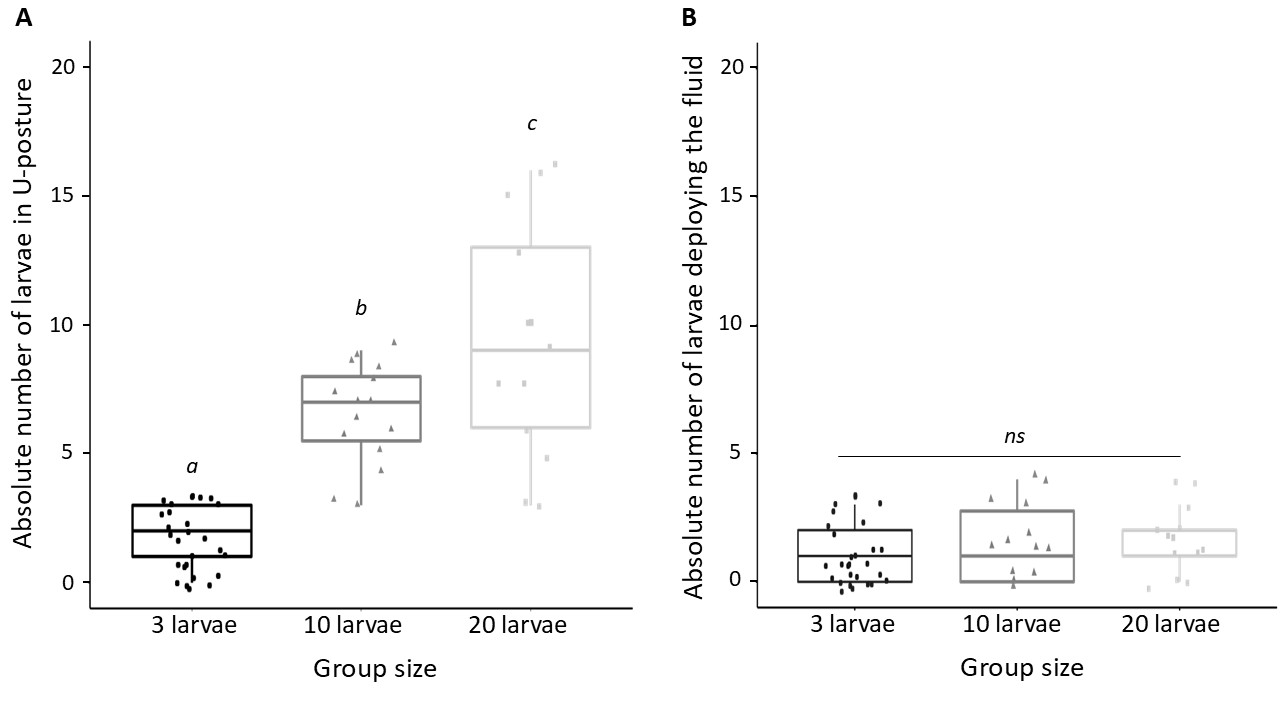
**Supplemental figures**

**Figure S1.** Effect of the group size on collective defence. (A) The absolute number of larvae exhibiting the defensive U-posture after a simulated attack is lower in groups of 3 larvae than in groups of 10 larvae and groups of 20 larvae. (B) The absolute number of larvae deploying the defensive fluid after a simulated attack is similar in all the group size. The box plots represent the median and interquartile range, with whiskers extending to 1.5 times the interquartile range, each point represents one individual. ns = no significant difference between treatments. For pairwise comparisons, different letters correspond to p < 0.05.

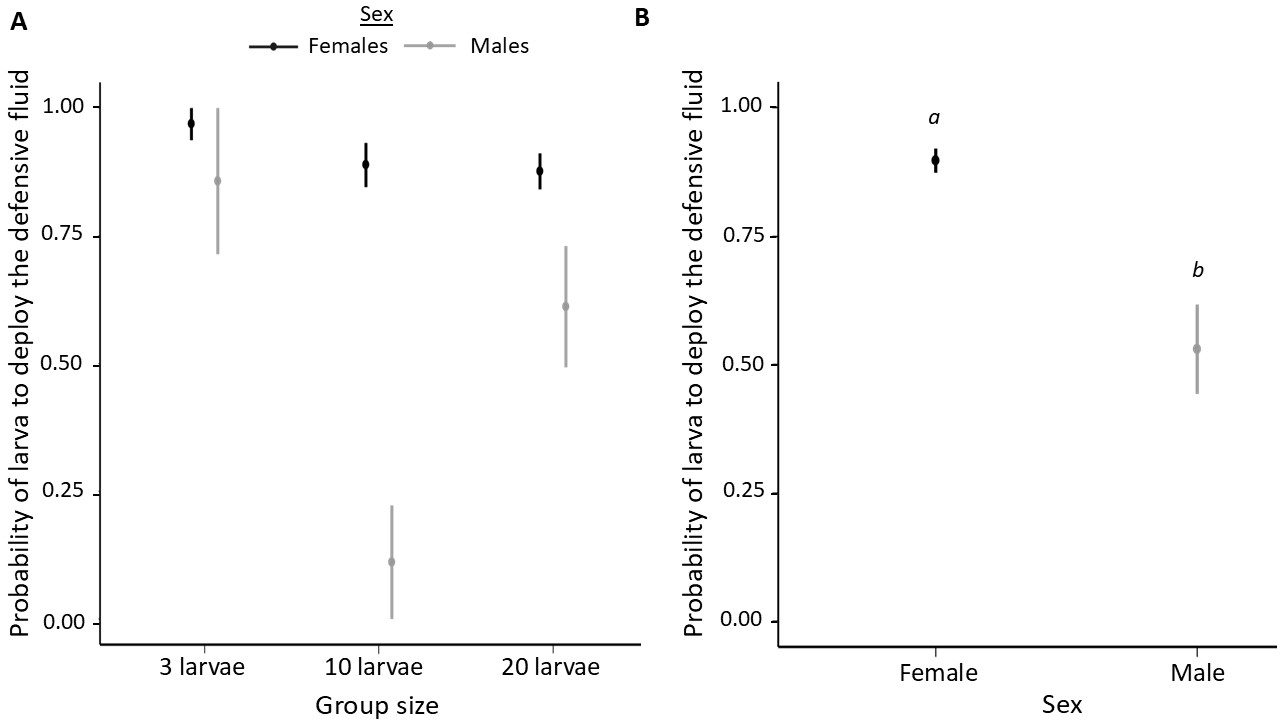

**Figure S2**. Effect of larval sex on their willingness to defend themselves. (A) Defensive fluid deployment was lower in groups of 10 larvae than in groups of 3 larvae but did not differ from groups of 20 larvae. (B) Females are more likely to deploy the defensive fluid than males. For pairwise comparisons, different letters indicate contrasts for which the 95% credible intervals do not overlap with zero.

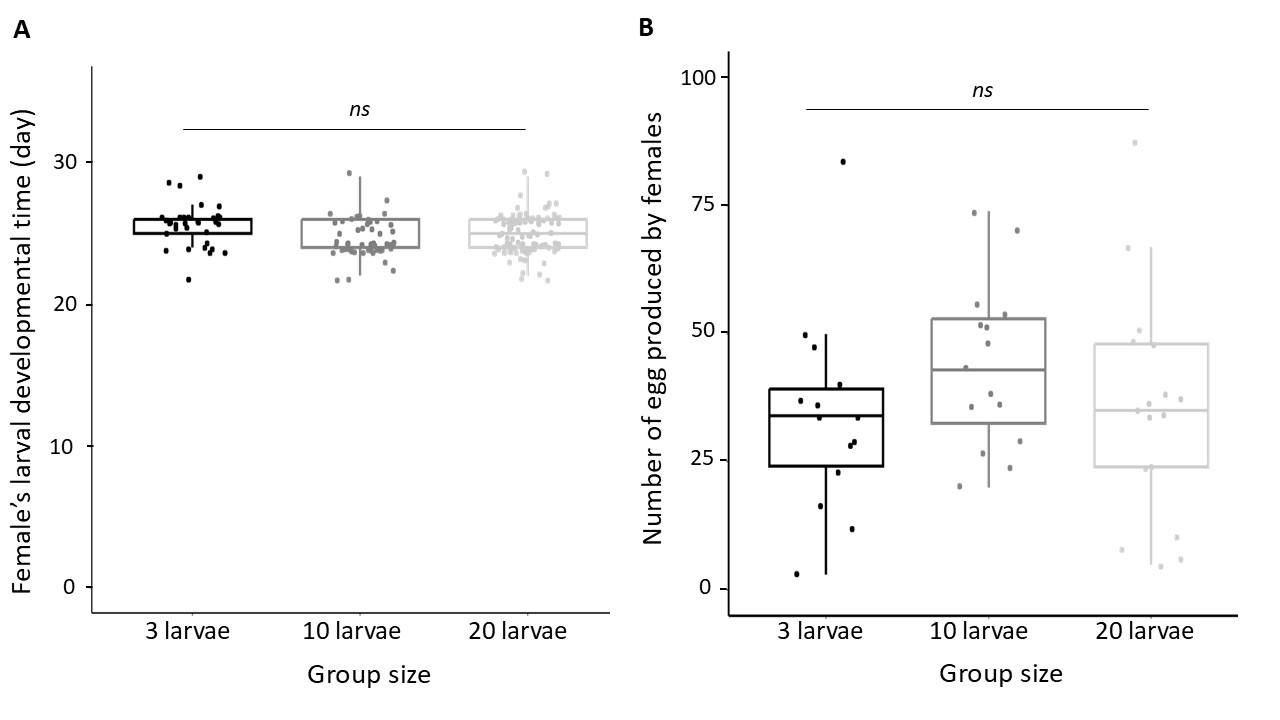

**Figure S3**. Effect of group size on females’ life history traits. (A) Female larvae raised in group of 3, 10 and 20 larvae develop equally fast, and (B) lay a similar number of eggs irrespective of the natal group size. The box plots represent the median and interquartile range, with whiskers extending to 1.5 times the interquartile range, each point represents one individual. ns = no significant difference between treatments.
